# CRISPRi-Seq for the Identification and Characterisation of Essential Mycobacterial Genes and Transcriptional Units

**DOI:** 10.1101/358275

**Authors:** Timothy J. de Wet, Irene Gobe, Musa M. Mhlanga, Digby F. Warner

## Abstract

High-throughput essentiality screens have enabled genome-wide assessments of the genetic requirements for growth and survival of a variety of bacteria in different experimental models. The reliance in many of these studies on transposon (Tn)-based gene inactivation has, however, limited the ability to probe essential gene function or design targeted screens. We interrogated the potential of targeted, large-scale, pooled CRISPR interference (CRISPRi)-based screens to extend conventional Tn approaches in mycobacteria through the capacity for positionally regulable gene repression. Here, we report the utility of the “CRISPRi-Seq” method for targeted, pooled essentiality screening, confirming strong overlap with Tn-Seq datasets. In addition, we exploit this high-throughput approach to provide insight into CRISPRi functionality. By interrogating polar effects and combining image-based phenotyping with CRISPRi-mediated depletion of selected essential genes, we demonstrate that CRISPRi-Seq can functionally validate Transcriptional Units within operons. Together, these observations suggest the utility of CRISPRi-Seq to provide insights into (myco)bacterial gene regulation and expression on a genome-wide scale.

## Introduction

Gene essentiality has been invoked extensively in both fundamental and applied studies of bacterial physiology and function^1^. For bacterial pathogens, classifications of essential genes can provide key insights into mechanisms of pathogenesis, as well as identifying potential vulnerabilities for new antibiotic drug-discovery^2^. It is not surprising, therefore, that elucidating essential genes and gene functions – whether as core or conditionally essential – has been a major focus in studies of *Mycobacterium tuberculosis*, causative agent of tuberculosis (TB). As for many other bacterial systems, genome-wide approaches to assigning gene essentiality in *M. tuberculosis* have primarily utilised forward genetic screens based on transposon (Tn) insertion mutagenesis^3^. One of the earliest applications of this technique, dubbed TraSH, for *tra*nsposon *s*ite *h*ybridization^4^, was quickly adapted in subsequent essentiality screens that incorporated NGS for transposon detection and analysis^5–7^, offering increasing depth of coverage and confidence in essentiality calls. This next generation of Tn-Seq approaches has been applied to questions of conditional essentiality in infection models^8^ and, more recently, to elucidate differential gene essentialities between clinical *M. tuberculosis* isolates^9^.

The indubitable utility of Tn mutagenesis in identifying essential genes contrasts with its inability to allow downstream analysis of the phenotypes of those very same essential genes: the insertions are, by definition, lethal and the mutants are eliminated from the pool^10^. To address this limitation, regulable knockdown systems incorporating inducible promoter replacements^11^, in some cases in combination with targeted protein degradation^12^, have been developed and enable controlled validation and nuanced phenotyping of essential and conditionally essential genes identified by Tn-Seq. These are difficult to scale, though, and complex to design, generate and validate, requiring labour-intensive maintenance and propagation. The pioneering development^13^ of a mycobacterial knockdown system based on CRISPR interference (CRISPRi) (Figure 1A) presents an efficient alternative for large-scale library construction, with the capacity for rapid validation and the potential to titrate repression^14^. However, since CRISPRi depends on the use of custom single guide RNAs (sgRNAs) which can have variable and unpredictable efficacies, and given the paucity of guidelines for the design of sgRNAs for variant Cas9 endonucleases^15^, initial characterization of the ability of a specific guide sequence to repress transcription represents a key requirement for the subsequent use of CRISPRi in phenotypic assays.

**Figure 1.**
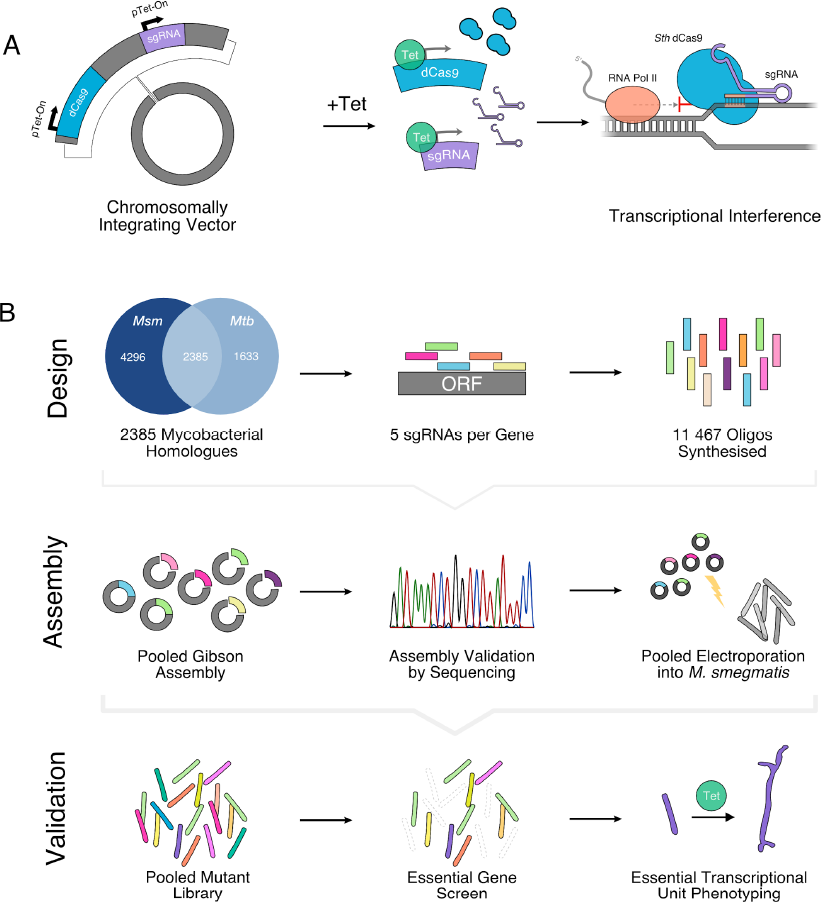
**Designing a Pooled CRISPRi Screen for Mycobacteria. (A)** The mycobacterial CRISPRi system^36^ drives production of dCas9 and a sgRNA from a Tetracycline-inducible promoter, off a single integrating plasmid. Addition of anhydrotetracycline (ATc) leads to transcriptional interference through steric hindrance of RNA polymerase. **(B)** A large-scale pooled library of sgRNAs was designed *in silico* to target *M. smegmatis (Msm)* homologues of *M. tuberculosis (Mtb)* genes. 11,467 sgRNAs, targeting 2,385 genes with up to five sgRNAs per gene, were produced as an oligo pool, and cloned into the delivery vector. The plasmid library was electroporated into *M. smegmatis,* creating a pooled, inducible knockdown library, which was used for a large-scale essentiality screen and subsequent downstream analysis.

CRISPR-based techniques have transformed eukaryotic cell biology, with large-scale pooled and arrayed CRISPRi libraries enabling high-throughput screens utilizing gain-of-function or loss-of-function assays to acquire insights into multiple biological mechanisms including cellular function, drug resistance and vulnerability^16^. More recently, the same approaches have gained traction in bacterial studies^17^, heralding the widespread application of this technology and derivatives to elucidate fundamental mechanisms of bacterial gene expression, regulation, and function^18–20^. As pooled libraries can incorporate guide redundancy to ensure efficient repression, pooled CRISPRi screens offer the opportunity to extend and supplement Tn-Seq, leveraging the power of scale to identify essential genes while empirically identifying the sgRNAs that are most effective at targeting them. In this way, CRISPRi screens can combine essentiality screening with rapid hit validation.

New tools require thorough characterisation. Whereas *E. coli* screens have utilised the catalytically inactivated dCas9 from *S. pyogenes (Spy)*^17^, the mycobacterial CRISPRi system incorporates dCas9 from *S. thermophilus (Sth)*^21^. It is necessary, therefore, to elucidate generalizable features that make for effective sgRNA design in this system, in addition to providing key insights into the factors which might impact gene essentiality classifications based on mycobacterial CRISPRi analyses. For example, uncertainty surrounds the vulnerability of CRISPRi to mis-assignments owing to polar effects: the simultaneous (untargeted) repression of genes downstream of a targeting sgRNA owing to transcriptional interference in the synthesis of a polycistronic mRNA. To date, characterisation of polar effects has depended primarily on the concept of the operon – a series of genes, often functionally related, that is transcribed in a single mRNA molecule. Little attention, though, has been paid to the observation that operons are described to fluctuate in structure: multiple mRNAs, or Transcriptional Units (TUs) can be produced from a single operon in a growth condition-dependent manner owing to the presence of alternative promoters, terminators, or other regulatory features (*e.g*., riboswitches)^22,23^. The potential susceptibility of CRISPRi to polar effects may, therefore, provide an opportunity to exploit pooled CRISPRi screens to explore operon biology at a genome-wide scale, using gene essentiality to identify and deconvolute functionally important TUs.

Here, we report the design, construction, and analysis of a pooled CRISPRi library in mycobacteria, specifically targeting *M. smegmatis* homologues of *M. tuberculosis* genes. To validate this strategy, termed CRISPRi-Seq, we compare our results to gene essentialities inferred from Tn-Seq analysis of *M. smegmatis*, benchmarked against Tn-Seq datasets for *M. tuberculosis*. We utilise CRISPRi-Seq to derive critical information about the characteristics making for effective targeting by an sgRNA, and empirically identify highly active sgRNAs for targeting essential genes. Finally, we leverage CRISPRi-Seq to probe operon biology, using polar effects to identify functionally-relevant TUs within operons, both on a genome-wide scale, and for a single operon containing essential genes for cell division and cell wall maintenance.

## Results

### A CRISPRi library targeting M. smegmatis homologues of M. tuberculosis genes

To provide a comparator for CRISPRi-Seq analyses of inferred gene essentiality, we performed Tn-Seq on wild-type *M. smegmatis* mc^2^155 after selection under standard aerobic growth conditions *in vitro*. The resulting data were analyzed using the Transit package^24^ and constitute the first published Tn-Seq essentiality analysis in *M. smegmatis* (Supplementary Data). For CRISPRi-Seq, we implemented a workflow incorporating *in silico* design of sgRNAs, pooled oligonucleotide synthesis and *en masse* plasmid delivery into a defined mycobacterial reporter strain (Figure 1B)^16^. In validating this approach, we targeted genes with identified homology^25^ between *M. smegmatis* and *M. tuberculosis* (Figure 1B), thereby focusing our screen on common mycobacterial functions relevant to the pathogen. According to our Tn-Seq analysis, the set of *M. tuberculosis* homologs comprised a combination of essential (ES) and non-essential (NE) genes, as well as genes affecting the *in vitro* growth rate – either growth defective (GD) or growth advantageous (GA) (Figure 2A). Of the 2,385 genes identified, 22 could not be targeted by Tn-Seq as they did not contain any -TA- dinucleotides required for random insertion of the MycoMarT7 phage^26^.

**Figure 2.**
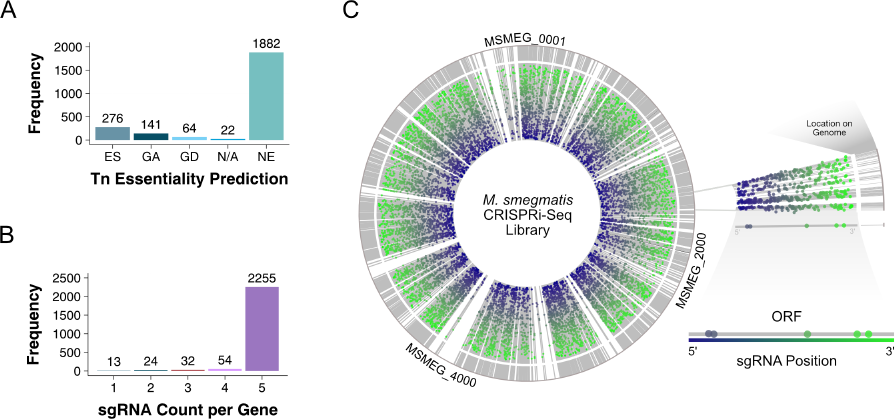
**A Pooled CRISPRi Library Targeting** ***M. smegmatis*** **Homologues of** ***M. tuberculosis*** **Genes. (A)** Tn-Seq analysis predicted that 276 of the selected genes were essential (ES), 205 affected growth rates (GA, Growth Advantaged; GD, Growth Defective) and 1,882 were non-essential (NE). The remaining 22 genes could not be targeted by Tn analysis (N/A) owing to their lack of TA sites. **(B)** A maximum of five sgRNAs were designed to target each gene to increase likelihood of effective knockdown. **(C)** The CRISPRi-Seq library targeted 2,385 genes, with approximately five sgRNAs targeting various points along each gene. Each gene is represented by a grey line, positioned by its accession number, in a circular coordinate system. Each sgRNA is represented by a single dot, and is positioned according to its relative position along the targeted gene, with 5’ sgRNAs in blue, and 3’ sgRNAs in green.

All sgRNAs were targeted to the non-template strand of the Open Reading Frame (ORF), defined as the distance between a bioinformatically identified start and stop codon^27^, to block transcription elongation. Optimal sgRNA selection required a custom bioinformatics pipeline (Supplementary Figure 1) that aimed to avoid off-target effects and potential constraints of oligonucleotide synthesis. From the resulting curated list, we selected up to five sgRNAs per gene (Figure 2B) to account for variability in guide efficacy. The sgRNA library, which comprised 11,467 sgRNAs across 2,385 genes (Figure 2C), was synthesised as a pool and cloned into the dCas9-expressing vector^21^ using modified Gibson assembly. Sequencing of the resulting plasmid library revealed that 99.99% of the sgRNAs were present, with only ten of the designed sgRNAs missing (Supplementary Figure 2).

### CRISPRi-Seq refines M. smegmatis gene essentiality predictions

To demonstrate the utility of CRISPRi-Seq in mycobacteria, we aimed to perform an essentiality screen in *M. smegmatis* (Figure 2A), validate against our Tn-Seq results, and generate essentiality calls on genes that could not be targeted by Tn-Seq. The pooled library was electroporated into a mutant *M. smegmatis* strain expressing a translational fusion of the chromosomal partitioning protein, ParB, to the mCherry fluorophore^28^. This ParB-mCherry reporter mutant was chosen as host strain as we considered DNA replication a useful indicator of cellular viability and function. After plating on standard 7H10 agar with or without anhydrotetracycline (ATc) – to induce or not induce CRISPRi, respectively^13^ – cells were scraped and sgRNAs sequenced from both pools. Analysis with MAGeCK-VISPR^29^, a quality control, analysis and visualisation package for CRISPR/Cas9 screens, revealed greater than 200-fold coverage per sgRNA for each condition (Supplementary Figure 3A). Notably, both Gini index analysis – a marker of uneven sgRNA distributions – and missing sgRNA counts increased with CRISPRi induction (Supplementary Figures 3B and 3C), supporting effective essential gene suppression. Moreover, the uninduced library overlapped closely with the pure plasmid library (Supplementary Figure 3D), suggesting minimal loss of coverage following electroporation into *M. smegmatis* and selection of transformants.

**Figure 3.**
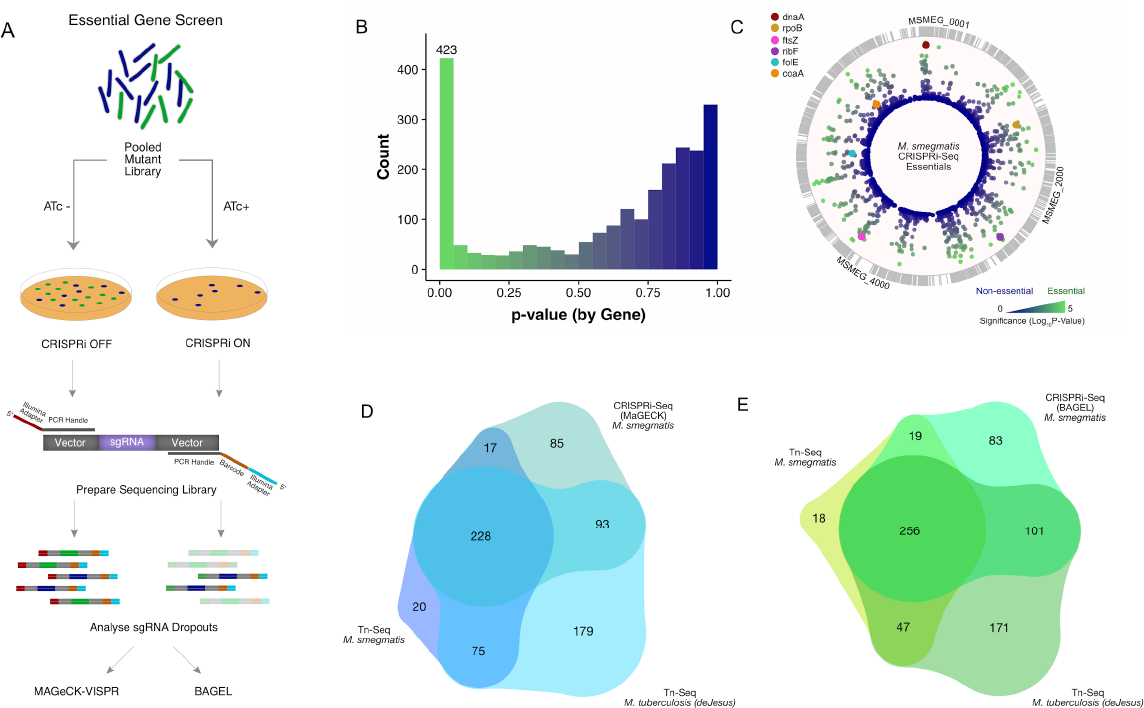
**– CRISPRi-Seq Identifies Essential** ***M. smegmatis*** **Genes. (A)** A pooled negative selection screen of the *M. smegmatis* CRISPRi library was performed. For selection, the pooled mutant library was plated on 7H10 agar with or without CRISPRi-inducing ATc. Colonies were scraped from their respective plates, genomic DNA prepped, and a targeted Illumina sequencing library of the sgRNA insertions prepared. **(B)** Analysis with MaGECK-VISPR, a quality-control, analysis and visualization package for CRISPR screens, identified 423 genes as significantly (*p* < 0.05) underrepresented, or essential, including **(C)** known essentials in cell division (*ftsZ)*, transcription (*rpoB*), DNA replication (*dnaA*) and metabolism (*ribF*, *folE*, *coaA*). **(D)** Comparison of CRISPRi analyses using MaGECK-VISPR to Tn-Seq of *M. tuberculosis* and *M. smegmatis* identified 228 universal essentials, which increased (E) to 256 (of 450 identified essentials) using the alternative analysis package, Bayesian Analysis of Gene Essentiality (BAGEL), trained with universal essentials and non-essentials.

Applying MAGeCK-VISPR with a Robust Ranking Algorithm identified 423 genes with significant underrepresentation on induction (Figure 3B). Closer analysis (Figure 3C) confirmed that genes encoding core functions in cell division (*ftsZ*^30^), transcription (*rpoB*^31^), and DNA replication (*dnaA*^31^) were identified as essential, whereas known non-essential processes, such as components of the mycobacterial SOS response (*e.g., dnaE2*^32^), were not. To confirm the inferred validity of the CRISPRi-Seq method, we compared the CRISPRi-derived essentiality calls with our own *M. smegmatis* Tn-Seq dataset (Figure 3D), as well as the recent saturating Tn-Seq dataset from *M. tuberculosis*^7^. For our comparisons, we considered all genes required for optimal growth – that is, those classified by Transit as “growth defect” (GD) or “essential” (ES) – as equivalent to an essentiality call by MAGeCK-VISPR. This analysis revealed significant overlap (228 genes) between the three datasets, with CRISPRi identifying a number of essential genes that had not been identified in our Tn-Seq screen.

Essentiality analysis is dependent on the bioinformatics pipeline utilised; therefore, we aimed to optimise our essentiality calls further by applying a Bayesian Analysis of Gene Essentiality (BAGEL) approach^33^ and training with genes identified by both MAGeCK and Tn-Seq as either essential (ES and GD by Tn) or non-essential (NE). BAGEL increased the number of essential gene calls to 459, and increased the extent of overlap between the datasets by 28 genes (Figure 3E), as well as increasing the number of genes not identified by our Tn-Seq dataset. Of the 22 genes excluded from the Tn-Seq analysis owing to their complete lack of -TA- sites, seven were identified as essential by BAGEL analysis, thereby highlighting the capacity of CRISPRi-Seq to provide essentiality data for genes deficient in the Tn insertion site. In addition, *M. smegmatis* mc^2^155 possesses a chromosomal duplication^34^ that is not present in *M. tuberculosis* and cannot be accurately classified by Tn-Seq; we noted that CRISPRi-Seq identified four essential genes within the duplicated region, insertions in which had been identified as conferring a growth advantage (GA) in the Tn-Seq analysis. Satisfyingly, Tn-Seq results from *M. tuberculosis*, where these genes are present in single copy, echoed the essentiality calls of CRISPRi-Seq.

### Knockdown efficacy depends on PAM Sequence, and is largely unaffected by sgRNA placement in the Open Reading Frame

In addition to assignments of essentiality, we aimed to utilise our dataset to quantify knockdown efficacies for sgRNAs targeting identified essential genes (Supplementary Data) and to gain insight into the factors which determine sgRNA efficacy.

The *Sth* dCas9 utilised in the mycobacterial CRISPRi system^21^ requires a 7 nucleotide protospacer adjacent motif (PAM) sequence whereas the better characterised *Spy* dCas9 recognises a 3 nucleotide PAM^35^. The longer *Sth* dCas9 sequence is necessary for CRISPR function and, although variations in the PAM sequence can be tolerated, they are associated with a decline in repression efficacy^21^. We examined the genome-scale impact of PAM sequence variation by analysing the relationship between the PAM Score – a measure of variations in the PAM sequence^36^ – and knockdown efficacy (Supplementary Table 1). For this, we focused on the sgRNAs targeting genes identified as essential by BAGEL. These 2,195 sgRNAs should result in a quantifiable decrease in read count following CRISPRi induction – correlating with effective repression and a decline in mycobacterial viability. For each sgRNA, we calculated the log2 ratio of read counts before and after induction of CRISPRi as a marker of knockdown efficacy. As noted previously^21^, lower PAM scores produced more effective knockdown (Figure 4A). Furthermore, when looking at the median values for each PAM score, a strong linear correlation (r = 0.855, *p* = 2.28e-6, Spearman’s Two-Sided) was observed (Figure 4A), confirming the importance of the PAM sequence in designing an effective sgRNA.

**Figure 4.**
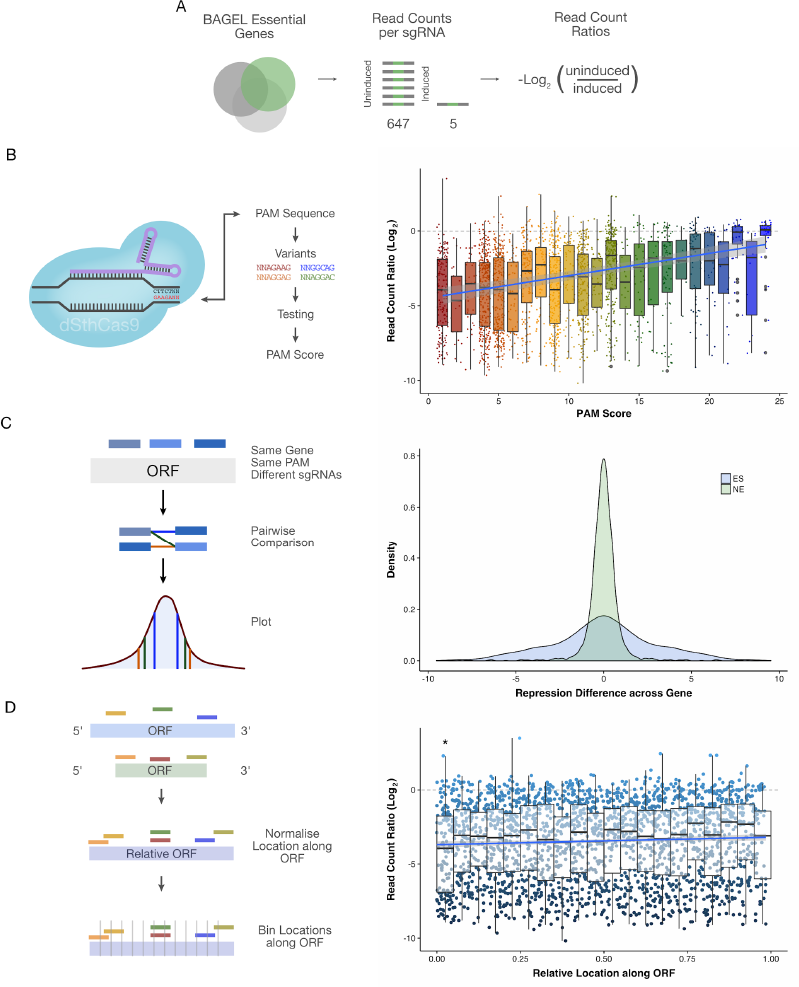
**PAM Score is the Primary Predictor of sgRNA Efficacy. (A)** Read Count Ratios, the log2 of the ratio of uninduced to induced read counts, were calculated for the 2,195 sgRNAs targeting essential genes, as defined by BAGEL analysis. (**B**) In general, lower PAM scores led to greater knockdown efficiency, although a range of efficacies was observed within each score. The Median Read Count Ratio of PAM Score classes showed a strong linear relationship (r = 0.855, *p* = 2.28e-6, Two-Sided Spearman’s) between PAM Score and Knockdown efficiency. (**C**) For sgRNAs with the same PAM Score, and targeting the same essential gene, pairwise comparisons of Read Count Ratios showed more variability than sgRNAs targeting non-essential genes, where the resultant variability was purely due to sampling stochasticity (*p* < 2.2e-16, Two Sided F-test). (**D**) No correlation was observed between sgRNA placement along the ORF and knockdown efficiency (r = 0.055, p = 0.009, Two-Sided Spearman’s). When binned into increments of 5%, median knockdown efficacy was significantly improved in the first 5% of the ORF, compared to random placement on the ORF (*p* = 0.0068, Two-Sided Wilcoxon Rank Sum test, 95% CI [−1.22, −0.19]).

Transcriptional repression by dCas9 depends on two probabilities^14^: the probability of the catalytically inactivated dCas9 nuclease blocking RNA polymerase on collision, and the probability of dCas9 occupying the target – itself a product of the dCas9 binding affinity, protein concentration, and rate of unbinding (either transcription-dependent or transcription-independent). Our data represent constant concentrations of dCas9 across the population and, for a single gene, presumably consistent rates of transcription-dependent collision. Therefore, we wondered if, for a single gene, PAM score alone determined dCas9 binding and unbinding and, consequently, could predict knockdown efficacy. For single genes, we calculated the differences in repression for sgRNAs with the same PAM scores (Figure 4C): if PAM score alone predicted efficacy, differences in repression should display a tight normal distribution. The distribution of differences across non-essential genes was only affected by sampling randomness (as there was no repression), and was more tightly distributed than the equivalent distribution for essential genes. This result suggested strongly that, though highly predictive of sgRNA efficacy, PAM score alone could not account for dCas9 binding and transcription-independent unbinding.

We also considered whether the location of the sgRNA on the ORF impacted sgRNA efficacy. Intuitively, we assumed that sgRNAs closer to the 5’ end of the ORF would lead to more effective knockdown. Strikingly, this was not the case (Figure 4C) with no clear relationship discernible (r = 0.055, *p* = 0.009, Two-Sided Spearman’s) between location on the ORF and sgRNA efficacy. However, we did observe that sgRNAs targeting the 5’-terminal 5% of the ORF were more effective(*p* = 0.0068, Two-Sided Wilcoxon Rank Sum test, 95% CI [−1.22, −0.19]), echoing a previous finding in *E. coli*^37^.

### CRISPRi-Seq identifies essential Transcriptional Units

Our CRISPRi-Seq analysis identified 83 *M. smegmatis* genes as essential which were not similarly classified by Tn-Seq in either the recent *M. tuberculosis* dataset^7^ or our own *M. smegmatis* dataset. As noted above, five of the 22 genes lacking -TA- sites were identified as essential by CRISPRi-Seq, representing one (predictable) benefit of CRISPRi-Seq over Tn-Seq. An additional 7 genes, when compared to either Sassetti^4^ or Griffin^5^ Tn screens, showed some evidence of essentiality. CRISPRi is known to have polar effects on downstream genes within operons^17,21,38^, potentially confounding calls of essentiality. Therefore, we asked if polar effects (Figure 5A) might account – even partially – for the 71 remaining differences in essentiality calls between CRISPRi-Seq and Tn-Seq.

**Figure 5.**
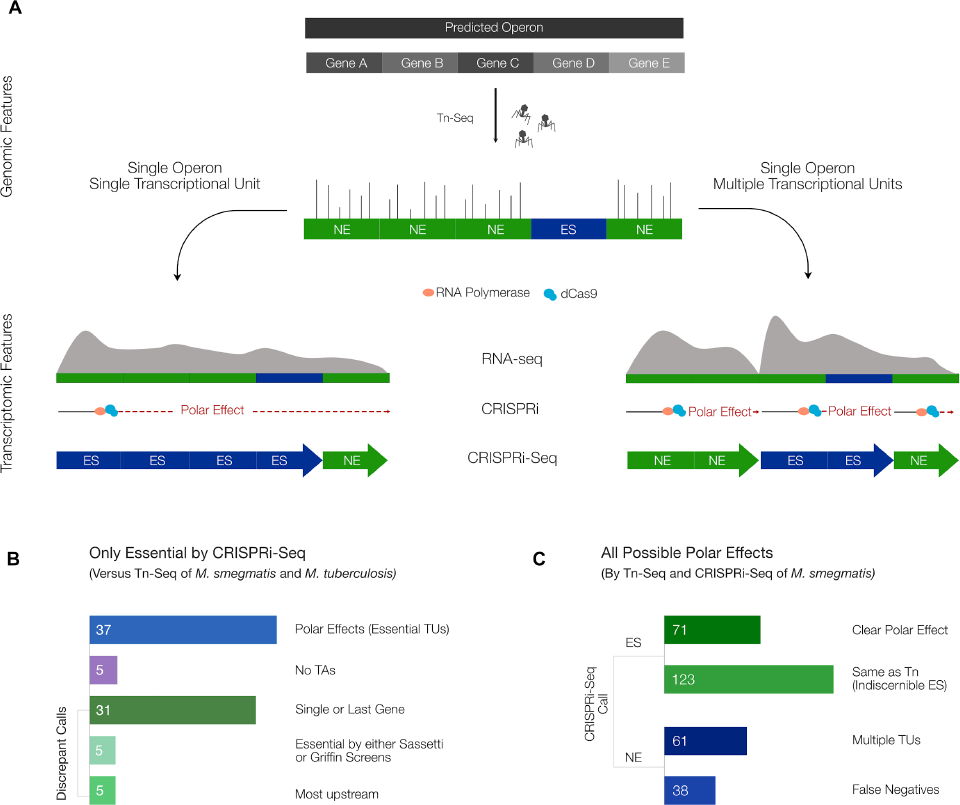
**CRISPRi-Seq Identifies Essential Transcriptional Units.** (**A**) An operon classically comprises a cluster of contiguous genes that are co-transcribed on a single mRNA transcript. However, operons can sometimes be transcribed as smaller, sub-operonic transcriptional units (TUs). Whereas Tn-Seq allows for analysis of single-gene essentiality within an operon. CRISPRi knocks-down an entire TU, and thus can obscure single-gene essentiality calls owing to polar effects. Comparing Tn-Seq calls with CRISPRi-Seq can therefore reveal the essentiality of a TU. (**B**) Eighty-three genes were called essential solely by CRISPRi-Seq. Five contained no TA sites and could not be targeted by Tn-Seq. Of the remainder, 31 were either not in an operon or were the last gene in a predicted operon; 5 were called essential by either Sassetti^4^ or Griffin^5^ Tn screens; and 5 were the most upstream gene in a predicted operon containing no downstream essentials. Together, these 41 genes represent discrepant calls between Tn-Seq and CRISPRi-Seq. The remaining 37 (45%) calls, though, had downstream essential genes in a predicted operon, thus revealing essential TUs. (**C**) There were 293 genes with possible operon-level polar effects – genes with downstream essential in a predicted operon. Seventy-one of these were clear polar effects, and were called ES solely in the CRISPRi-Seq analysis. The other 123 were ES by both Tn-Seq and CRISPRi-Seq, and therefore could not be distinguished as essential genes or TUs. There were 99 genes which did not exhibit the expected polar effects. Analysis of publicly available RNA-Seq data (smegmatis.wadsworth.org) revealed that 61 (62%) of the “missing” polar effects occurred where operons consisted of multiple TUs (as suggested by the RNA-Seq trace), thereby negating polar effects on downstream essential genes. The remaining 38 genes represented discrepant calls.

Computational predictions of operons have been performed in a large number of bacterial species^39^. However, these predictions largely depend on a definition of the operon that excludes overlap between operons and ignores variant transcripts within an operon^23^. For initial polar effect analysis, we adopted non-contiguous operon prediction by DOOR as standard. We excluded 31 of the 71 discrepant calls as they were predicted to be either standalone genes or the final gene in a predicted operon. These discrepant calls may represent false-negatives of the Tn-Seq analysis, or false positives from CRISPRi-Seq. Notably, of the remaining 40 genes, 37 (52% of the 71 initial genes) were associated with downstream genes (in the same predicted operon) described as essential by either Tn-Seq or CRISPRi-Seq.

We were intrigued by the impact of polar effects on our CRISPR-Seq results, and sought to understand how this phenomenon affected the validity of essentiality calls more generally. As upstream polar effects are infrequently observed^17^, we examined all downstream polar effects: that is, we searched for all instances in which a gene was located upstream of an essential gene (as classified by Tn-Seq or CRISPRi-Seq) in a computationally predicted operon^39^(Figure 5A). There were 293 genes located upstream of essential genes in predicted operons, of which 194 were called ES by CRISPRi-Seq. For 123 of these, Tn-Seq also returned a classification of ES, highlighting a key limitation in the CRISPRi-Seq methodology: only the most downstream gene in an operon can be called essential with certainty. The remaining 71 represent clear polar effects.

A large number of genes (99) with possible polar effects was classified NE by CRISPRi-Seq. This was unexpected, and prompted further analysis of publicly available RNA-Seq datasets^40^. Although operons have traditionally been considered a fixed means of co-regulating genes, evidence exists that transcription across an operon is more dynamic than initially appreciated: whole operons, or portions of operons, can be flexibly transcribed in a growth condition-dependent manner^22^, perhaps consistent with the dynamism of higher order chromosomal organization^41,42^. In RNA-Seq data, TUs are observable with peaks in the read counts at the position of transcription start sites (TSSs), followed by consistent transcription (as determined by read counts) of downstream genes, and eventual transcription termination^43^ (Figure 5A). Strikingly, of the 99 genes whose depletion was not associated with polar effects, the majority (62%) appeared to be located in TUs that differed from their predicted operon structures: that is, these genes were characterized by an increase in read count *prior* to the downstream essential gene (Supplementary data). In turn, this suggested that discrepancies between Tn-Seq and CRISPRi-Seq might be used in conjunction with RNASeq data and information about TSS locations to identify functional TUs.

### CRISPRi-Seq enables phenotyping and functional validation of Transcriptional Units

While systems-level differences between Tn-Seq and CRISPRi-Seq calls can be used to identify essential TUs, difficulties arise where Tn-Seq and CRISPRi-Seq converge on calls of essentiality: in an operon where all calls are essential, discrepancies cannot be used to identify polar effects, nor discern functional TUs, even where RNA-Seq traces vary within the predicted operon. We speculated that the ability to phenotype predicted essential genes rapidly – for example, using image-based analyses – might be leveraged to validate functional TUs within putative operons entirely comprising essential genes. The division and cell wall (*dcw*) operon presented an intriguing system to evaluate this idea, offering an opportunity to combine transcriptomic data, CRISPRi-induced polar effects, and terminal image-based phenotyping to validate the functional TUs within this operon. Four genes (*murC, murD, mraY* and *murG*) within the *dcw* operon are involved in peptidoglycan precursor synthesis (Figure 6A), and are interspersed with the cell-division genes, *ftsW* and *ftsQ.*

**Figure 6.**
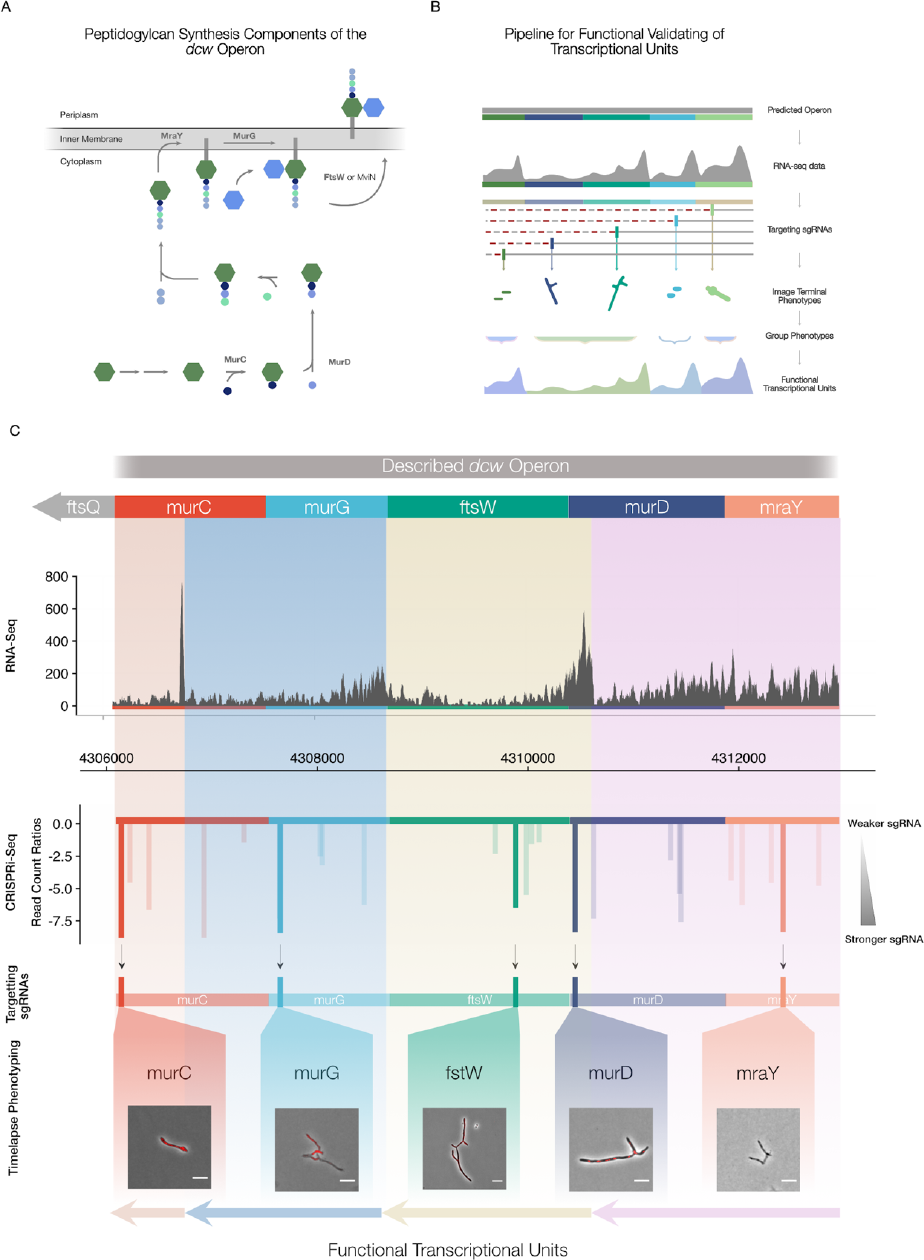
**CRISPRi-Seq Allows for Phenotyping and Functional Validation of Transcriptional Units.** (**A**) Synthesis of the peptidoglycan precursor, lipid II, requires the expression of a gene cluster comprising MurC, MurD, MraY, MurG and FtsW. (**B**) We predicted RNA-Seq, CRISPRi-Seq and terminal image-based phenotyping could be combined to phenotype various knockdowns along an operon, and determine functional TUs within the operon. (C) Using this pipeline, highly active sgRNAs were chosen to target each gene along the described *dcw* operon. Microscopy of terminal morphologies allowed grouping of phenotypes into functional TUs that corresponded to RNA-Seq read profiles.

Previous experimental observations support the classification of *dcw* as an operon in *Mtb*^44^, and this has also been suggested by RNA-Seq in *M. smegmatis*^45^. However, other publically available RNA-Seq data of *M. smegmatis* (smegmatis.wadsworth.org) indicate that there may be multiple TUs present within the operon (Figure 6C). To discern possible functional TUs within *dcw*, we targeted five genes situated centrally in the operon and identified as essential by both CRISPRi-Seq and Tn-Seq. For these, we re-synthesised the most active sgRNAs (Supplementary data) to create single-knockdown clones in the *parB-mCherry* reporter background. Importantly, we were able to select highly active sgRNAs located at different points across *dcw*, with the exception of *ftsQ*, for which no highly active sgRNAs had been identified in our screen. Following induction of CRISPRi in liquid culture, we phenotyped cells via microscopic imaging after 18 hours of exposure to ATc on solid agar pads. In addition, we performed time-lapse imaging of the mutants carrying knock-downs in the peptidoglycan biosynthesis components of the operon (Figure 6C; Supplementary movies).

If *dcw* functioned as a single TU under the conditions tested, sgRNAs targeting 5’ upstream of *ftsQ* and *ftsW* should have produced polar effects, thereby phenocopying disruption of the downstream genes – that is, targeted depletion of *mraY* or *murD* (upstream of *ftsQ*) or *murG* or *murC* (upstream of *ftsW*) should have resulted in the filamentation phenotype consistently reported in *fts* knockdowns^46^. However, terminal phenotyping demonstrated this not to be the case. Silencing of the most 5’ upstream gene in the operon, *mraY,* produced smaller cells that rapidly burst, consistent with impaired peptidoglycan maintenance (Figure 6C, Supplementary Movie 1 and Figure 4). In contrast, the *murD* sgRNA – targeted to the distal 3’ end of the *murD* ORF – triggered filamentation, phenocopying *ftsW* depletion (Figure 6C, Supplementary Movie 2 and Figure 4). Knock-downs of *murC* and *murG* also resulted in defects in cell wall structure and did not produce filamentation despite being upstream of *ftsQ* (Figure 6C, Supplementary Movie 3 and 4). Notably, by combining the locations of the targeting sgRNAs and their resultant time-lapse phenotypes with transcriptomic data, we observed that the phenotypes of particular sgRNAs grouped within transcriptional segments between potential TSSs, as indicated by upticks in the RNA-Seq trace. That is, the morphological phenotypes following targeted gene depletion provided strong evidence of underlying TU structure within the *dcw* operon. This result indicates that *dcw* does not act as a single functional unit in exponentially growing cells under standard laboratory conditions, in turn supporting the utility of CRISPRi-Seq for probing the functional architecture of (myco)bacterial operons.

## Discussion

Evidence that the bacterial genome is structurally and functionally dynamic^42,47–49^ supports the need for flexible, well characterised tools for high-throughput functional genomics. Here, we report the outcomes of a large-scale pooled knock-down screen in *M. smegmatis* utilising a recently developed CRISPRi system based on *Sth* dCas9^21^. In addition to validating CRISPRi-Seq for gene essentiality calling, our results offer key insights into the factors which determine effective application of this approach at scale, in turn revealing its potential utility as a tool for functional genomics and transcriptomics.

Our screen targeted coding regions of *M. smegmatis* genes with homologs in *M. tuberculosis.* This approach, which contrasted with recent genome-wide pooled CRISPRi screens in *E. coli* ^17^, facilitated the use of multiple sgRNAs for each targeted gene while providing the opportunity to leverage insight from *M. tuberculosis* systems biology. Comparing the CRISPRi-Seq essentiality calls to corresponding Tn-Seq data from *M. smegmatis* and published *M. tuberculosis* datasets^4,5,7^ validated the utility of this method for pooled essentiality screening in mycobacteria: CRISPRi-Seq identified 80% of genes classified as essential or growth defective by Tn-Seq in *M. smegmatis*, with the discrepant calls likely reflecting inherent limitations in either method. Closer analysis of these discrepancies suggests that CRISPRi-Seq has a false discovery rate of approximately 15.5% owing to polar effects. Moreover, a significant portion (50%) of the universally identified essential genes could not be definitively assigned as essential by CRISPRi-Seq alone owing to their operonic structure: essential genes are frequently grouped in operons, and CRISPRi-Seq can definitively identify only the operon, or final gene in the operon, as essential. These observations are similar to those reported recently in *E. coli*^50^.

For single-gene essentiality calling, Tn-Seq may remain the gold standard – even though unpredictable polar effects remain a potential confounder^51^. CRISPRi-Seq does, however, offer several significant benefits over Tn-Seq. For example, unlike Tn insertion mutagenesis, CRISPRi can simultaneously target ORFs located in a duplicated chromosomal region. In addition, CRISPRi-Seq libraries can be flexibly designed to focus on subsets of genes, potentially preventing bottle-necking sometimes observed in Tn-Seq^10^. Moreover, the inducible nature of CRISPRi-Seq suggests significant potential for novel screens based on phenotypes other than death^52–54^ as libraries can include essential genes, and offer the potential to titrate essential gene knockdown^14^. A first example of this type of screen was performed recently in *E. coli*^50^.

Recent work utilising the *Spy* dCas9 in *E. coli* has shown that repression efficiency can vary significantly across the same gene for different sgRNAs^17^. By focusing on identified *M. smegmatis* essential genes and using viability as a marker of guide efficacy, we observed the same effect with *Sth* dCas9 in the mycobacterial CRISPRi system. Here, the capacity of the *Sth* dCas9 to recognise variations in the PAM sequence, but with a negative impact on knockdown efficacies, was strongly evident. In general, it appeared that low PAM Scores resulted in more effective knockdown, although some sgRNAs with PAM Scores above the recommended value of 15 (Ref. 13) were effective. Importantly, PAM Score alone was not predictive of knockdown efficacy, even for sgRNAs targeting the same ORF; this reinforces the pervasive difficulty in sgRNA selection, while highlighting the value of the empirical guide efficacy data generated by CRISPRi-Seq. Similar to CRISPRi in *E. coli*^17^, we observed that location of the sgRNA sequence anywhere within the ORF might lead to effective knockdown; this was consistent with the report in mycobacteria^21^, but in stark contrast to early descriptions of bacterial CRISPRi^38^. Interestingly, as described elsewhere^37^, the increased resolution of CRISPRi-Seq established that the first 5% of the ORF appeared to produce more effective knockdown – possibly owing to the resulting proximity of the dCas9 complex to the promoter.

We investigated a target organism, *M. smegmatis*, with different (slower) growth and structural (Gram positive) characteristics from the *E. coli* workhorse utilised in other pooled CRISPRi screens^17^. In addition, the CRISPRi system employed a dCas9 of distinct origin. Perhaps owing to these differences, and the dominant effect of PAM Score, we were unable to identify the “bad seed” effect – off-target effects due to sgRNA seed similarity – described previously^17^. We attempted to leverage our dataset to design analysis pipelines that might increase our ability to select highly-active sgRNAs, as has been described for eukaryotic systems^55^, however these efforts have proved unsuccessful to date (scripts available on request). Libraries designed expressly to probe these aspects may provide more insight.

Although polar effects might be considered a confounder to the analysis of CRISPRi-Seq data, we combined operon predictions and the absence of predicted polar effects with publically available RNA-Seq data to gain fundamental insights into operon biology. The detection via this analysis of 71 clear polar effects provides experimental validation of the corresponding operon structures. More interestingly, the absence of polar effects in predicted operons led us to identify multiple TUs within 60% of these predicted operons. Attempts have been made at reclassifying operon structure using RNA-Seq data^45^; however, the functional TUs identified in our screen, when combined with an alternative RNA-Seq dataset, contradict a significant proportion of the reclassified operons (Supplementary Data). These discrepancies may be attributable to functional TUs changing in a condition-dependent manner^23,39^, or to the presence of multiple, overlapping TUs. Further experimental validation is required to clarify this possibility, preferably utilizing long-read RNA sequencing.

The results reported here demonstrate the utility of RNA-Seq for informing optimal guide placement and identifying potential polar effects. Analyzing discrepancies between *dcw* operon descriptions^29,44^ and RNASeq data on a single gene, single guide, single cell level, also revealed the potential of CRISPRi-Seq to probe the functional transcriptome – specifically, enabling the functional demonstration of multiple TUs within a single operon. The transcriptional uncoupling of these core components of peptidoglycan synthesis and cell division within a single operon contrasts with a traditional understanding of the operon as a dominant mediator of co-transcription. Moreover, given recent interest in the spatial and temporal coordination of cell wall construction and maintenance in mycobacteria^56–59^, the detection of TUs within *dcw* suggests further research is required to reveal the full suite of regulatory mechanisms employed in ensuring optimal function of this and related pathways. To this end, the identification of highly active sgRNAs at various points within *dcw* (and other operons) might be very useful, particularly when combined with the expanding suite of available chemical biology reagents and analytical techniques for probing mycobacterial cell structure and function^60–64^.

The notion of bacterial genomes as static has been discarded through the results of multiple post-genomic techniques which have revealed the capacity for dynamic alterations in, and interactions between, chromosomal structure^41,49^, composition, and function^42,47,48^. Even gene essentiality, long considered a binary trait, is now recognized as conditional and evolvable^1^, an insight enabled by – and in turn demanding – adaptable and flexible tools to investigate malleable genomic structures and functions. CRISPRi-Seq might offer a valuable new addition to the toolbox.

## Acknowledgements

We are very grateful to Mandy Mason, Anastasia Koch, and Valerie Mizrahi for helpful discussions and critical review of the manuscript, Marcus Gawronsky for assisting with the attempted machine-learning analysis of sgRNA efficacy, and Stephanie Fanucchi for help with figure design. Our thanks to Jeremy Rock and Sarah Fortune for their generous gift of the CRISPRi plasmids as well as technical advice on sgRNA design; thanks also to Chris Sassetti for the MycoMar transposon; and to Isabella Santi and John McKinney for the *M. smegmatis* ParB-mCherry reporter mutant. This work was funded by grants from the US National Institute of Child Health and Human Development (NICHD) U01HD085531 (to DFW); the Department of Science and Technology (DST) of South Africa; the South African Medical Research Council (SAMRC); the SAMRC SHIP initiative (to MMM); DST Centre of Competence Grant (to MMM); CSIR Parliamentary Grant (to MMM); and the University of Cape Town. MMM is a Chan Zuckerberg Investigator of the Chan Zuckerberg Initiative. We gratefully acknowledge the support of the SAMRC (Doctoral Scholarship to TdW), and the Research Council of Norway (INTPART, AMR-PART) for work on antimicrobial resistance in mycobacteria (to DFW).

## Author Contributions

Study conceptualization and design: TdW, MMM, DFW. *M. smegmatis* Tn-Seq: IG. CRISPRi-Seq library design, assembly, and analysis: TdW. *M. smegmatis* CRISPRi-Seq screen: TdW. Fluorescence and time-lapse imaging: TdW. Manuscript preparation: TdW, MMM, DFW.

## Data Availablity

The datasets generated during and/or analysed during the current study are available in the Open Science Framework repository, DOI 10.17605/OSF.IO/6X7VB. Additional data generated and/or analysed during this study are included in this published article (and its supplementary information files).

## Materials And Methods

### Construction of Transposon (Tn) Insertion Libraries

MycoMarT7 phage stock preparation and *M. smegmatis* transduction was performed as described^65^. A 200 ml culture of M. smegmatis WT strain was grown to an OD600 of 0.6 – 0.8, pelleted by spinning at 4000 RCF for 10 minutes, and the pellets washed three times with an equal volume of pre-warmed (37°C) MP buffer. Mycobacterial cells were resuspended in 4 ml of MP buffer. For transduction, 2 ml of phage stock was added to the cells and the culture was incubated at 37°C for 7 hours or overnight. After transduction, cells were pelleted and washed 3 times with PBS supplemented with 0.1% Tween80. Finally, the cells were resuspended in 6 ml of PBS-Tween80 and plated on 7H10 agar supplemented with 0.5% glycerol, 10% Middlebrook OADC enrichment (Difco), 0.1% Tween80 and 20 µg/ml kanamycin. Aliquots of 500 µl of cells were spread on each plate giving a total of 12 plates. Plates were incubated 37°C for 2 – 3 days after which cells were harvested by scraping and re-suspended in 7H9-OADC-Tween80 broth supplemented with 15% glycerol. The library was then frozen at −80°C in 1 ml aliquots.

### Genomic DNA Extraction and Illumina Library Preparation for Tn-Seq

Acoustic fragmentation was carried out using a Covaris M220 Focused-Ultrasonicator with the recommended protocol for a 300 bp target, as per manufacturer’s instructions. Random DNA shearing was followed by repair of DNA ends (End-It^TM^ DNA End-Repair Kit, Epicentre). A-tailing of fragments was carried out using dATPs and Taq DNA polymerase as described^65^. Finally, amplification of Tn–chromosomal junctions was performed as outlined^66^: Eight 30 µl reactions were performed for each library. The 8 reactions were then pooled and electrophoresed on a 2% TAE agarose gel. Fragments of sizes 250 – 600 bp were excised and gel purified for each library. After purification, a hemi-nested PCR was performed to add sequences required for Illumina sequencing. Samples were quantified by Qubit fluorometer.

### NGS of Tn libraries and Data Processing and Analysis

Prepared libraries of 3 biological replicates were sequenced on an Illumina MiSeq (Oklahoma Medical Research Foundation, OK, USA). At least five million 2 x 300 PE (Paired End) reads per sample were obtained. TPP Statistical analysis was carried out using TRANSIT (v2.1.2)^24^, and the default Hidden Markov Model method (HMM) used to generate essentiality calls. Default settings were used. In total, 21 119 053 reads were mapped to the mc^2^155 genome, achieving a density of 76.2%.

### CRISPRi Library Design

A list of all possible sgRNAs in the *M. smegmatis* mc^2^155 genome was obtained from Jeremy Rock. Manual curation narrowed down the list of genes to those with identified homologues between *M. smegmatis* and *M. tuberculosis*^25^. For each sgRNA in the ORF of the selected genes, we calculated the GC content, number of homopolymer repeats, and a “mismatch” score by aligning the sgRNA to the *M. smegmatis* genome using Rbowtie^67^. We proceeded to rank sgRNAs, in order, by PAM score, mismatch score, GC content and homopolymer repeat count. The top 5 ranked guides were selected for each gene, after which homologous vector sequence was added to either side of the sgRNA sequence to produce a single ˜109 bp sequence. The complete script is available at osf.io/zh5t9. 11,467 oligos (Supplementary data) were designed in this way, and synthesised by CustomArray (Bothell, WA).

### CRISPRi Plasmid Library Assembly

The assembly protocol was adapted from a previous report^16^. The vector backbone was digested overnight with BsmBI (NEB), and gel purified with a Monarch Gel Purification Kit (NEB). For assembly of the library, we amplified the ssDNA oligos with PCR primers targeting the constant vector sequence present on each oligo, and Q5 Hot-Start High-Fidelity Master Mix (NEB). The resultant dsDNA was gel purified with the aforementioned kit. A Gibson assembly was performed using NEBuilder^®^ HiFi DNA Assembly Cloning Kit (NEB), and the entire reaction concentrated using GlycoBlue Coprecipitant (Thermo). The concentrated reaction was electroporated into Endura Electrocompetent Cells (Lucigen), in three 50 µl reactions. Each individual reaction was plated onto low-salt LB plates containing Kanamycin (50 µg/ml) and incubated at 37°C overnight. Transformants were scraped, pooled and maxi prepped using a ZymoPure maxiprep kit (Zymo Research).

### CRISPRi Library Validation

The pooled plasmid library was sequenced through targeted amplicon sequencing of the variable sgRNA region. Briefly, custom Illumina primers targeted to the vector sequence surrounding the sgRNA were used to PCR amplify the region of interest, in a single PCR reaction, with Q5 Hot-Start High-Fidelity Master Mix (NEB). The reaction product was gel purified and quantified using a Nanodrop and Qubit, before being sequenced on an Illumina MiSeq (myGenomics Inc., GA, USA), providing ˜10 million PE reads. Reads were processed using an adapted version of the script described by Joung et al.^16^ calculating library coverage, and skew (the adapted script is available at osf.io/zh5t9).

### CRISPRi M. smegmatis Library Creation

The validated plasmid library was electroporated into an *M. smegmatis* mc^2^155 reporter mutant expressing a fusion of the ParB protein with mCherry from its native locus^28^. Successful transformants were selected on solid 7H10 agar supplemented with OADC and Glycerol (0.2%), containing Kanamycin (20µg/ml) following incubation at 37°C. Transformants were scraped into 20ml of 7H9 supplemented with OADC, 0.2% glycerol and 0.05% Tween80 with Kanamycin (20µg/ml), and were vortexed with glass beads, water-bath sonicated to break up clumps, and centrifuged briefly at 200 RCF to pellet remaining large clumps. The supernatant was removed, and aliquots created. Glycerol was added to a final concentration of 33% v/v and aliquots of the library were stored at −80°C. Prior to use, aliquots were filtered through a 5µm filter (Millipore) to ensure a single-cell suspension. The single cell suspensions were validated by microscopy prior to use, and the CFU counts determined.

### CRISPRi Negative Selection Screen

For the negative selection screen, equal volumes of ˜10^5^ CFU were plated on 7H10 agar plates containing Kanamycin (20µg/ml), with or without ATc (100ng/ml). Plates were incubated at 37°C for 2-3 days until visible colonies formed, after which the plate was scraped and the genomic DNA prepped using a standard CTAB extraction. Three replicates were performed for each selection screen. An amplicon library of the sgRNA sequences was prepared from the genomic DNA by PCR amplifying the vector sequence as described for plasmid library prep. Sequencing was performed by myGenomics Inc.(USA) on an Illumina MiSeq, providing 2 million PE reads for each replicate. Reads were pre-processed using cutadapt(v1.1.4)^68^ to remove primer sequence. To analyse data, MAGeCK-VISPR (v0.5.3)^29^ and BAGEL (v0.91)^33^ were used to generate calls of essentiality. Scripts and data used for analysis are available at osf.io/zh5t9.

### Polar Effect Analysis

For polar effect analysis, we obtained operon predictions from DOOR^39^ and combined these with our Tn-Seq essentiality data using a custom R script. For each gene in an operon, we noted whether or not there was an essential or growth defect-associated gene located downstream in that particular operon, we further noted if a gene was the only gene in an operon, or the last gene in an operon. To examine the presence of polar effects, we asked how many genes called non-essential by Tn-Seq of *M. smegmatis*, were essential by CRISPRi – those that had downstream essential genes (defined by either Tn-Seq or CRISPRi-Seq of *M. smegmatis)* in the same operon were considered polar effects. Genes that were called non-essential by Tn-Seq, and non-essential by CRISPRi, but had downstream essentials were manually examined by observation of RNA-Seq data (smegmatis.wadsworth.org). The full R analysis script and raw data are available at osf.io/xs7q2.

### sgRNA Efficacy Analysis

For sgRNA efficacy analysis, we selected all sgRNAs targeting genes identified as essential by BAGEL. For each sgRNA, we calculated a suppression ratio by taking the log2 of the ratio of induced to uninduced read counts. For analysis of PAM score effects, we compared the suppression ratios between PAM scores. To discern the impact of PAM on single genes, we calculated pairwise comparisons of sgRNA log2 ratios for sgRNAs targeting a single gene, with the same PAM score. We compared the variance of essential, to non-essential gene pairwise comparisons. For analysis of the impact on efficacy of location of an sgRNA within an ORF, we calculated the relative location along the ORF by dividing the distance of the sgRNA from the TSS by the total length of the ORF. The suppression ratio was plotted against this relative location. The full R analysis script and raw data are available at osf.io/8qd75.

### Phenotyping of TUs

We constructed single-gene knockdowns by synthesising short oligonucleotides corresponding to highly-active sgRNAs identified in our pooled screen. These were annealed and ligated into the same plasmid used for screening, as described previously^21^. Plasmids were electroporated into the same *M. smegmatis*ParB-mCherry strain used for the pooled screen. Strain knockdowns were induced, and imaged, as described below.

### Microscopy

For imaging CRISPRi terminal phenotypes of individual mutants, an exponential phase culture (OD600 ˜ 0.8) of each mutant was diluted 1:64 into fresh 7H9 OADC containing ATc (100ng/ml) and Kanamycin (20µg/ml), and incubated at 37°C for 18 hours with shaking. Cells were inoculated onto low-melt agarose pads and imaged on a Zeiss AxioObserver using a 100X, 1.4NA Objective with Phase Contrast and Colibri.7 fluorescent illumination system. Images were captured using a Zeiss Axiocam 503. For strains with terminal phenotypes appearing lytic, time-lapse microscopy was performed.

### Time-lapse Microscopy

For time-lapse microscopy, 1.5% low-melt agarose pads were prepared and were embedded with 7H9 media supplemented with OADC and glycerol. For induction of CRISPRi, 100ng/ml of ATc was added to the pad. Exponential phase cells (OD600 ˜ 0.8) were diluted to an OD of 0.1, with media containing 100ng/ml ATc and Kanamycin (20µg/ml), and exposed to ATc at 37°C for 6 hours. The exposed culture was filtered through a 5µm filter, and inoculated onto pads. Pads were imaged in the base of a glass-bottomed petri dish, on the Zeiss Axio Observer described previously, within an incubated stage at 37°C. Images were taken every 15 minutes, for approximately 18 hours.

